# Structural features of an infectious recombinant PrP^Sc^ prion using solid state NMR

**DOI:** 10.1101/2020.04.08.032839

**Authors:** Manuel Martín-Pastor, Yaiza B. Codeseira, Giovanni Spagnolli, Hasier Eraña, Leticia C. Fernández, Susana Bravo, Alba Iglesias, Rafael López-Moreno, Sonia Veiga, Emiliano Biasini, Víctor M. Sánchez-Pedregal, Joaquín Castilla, Jesús R. Requena

## Abstract

PrP^Sc^, the first described and most notorious prion, is the only protein known to cause epidemics of deadly disease. Its properties are encoded in its unique structure. Here we report a first solid state NMR study of a uniformly labelled (U-^13^C,^15^N)-Bank vole (BV) infectious recombinant PrP^Sc^ prion. C-C, C-H and N-H spectra were obtained with MAS rotation of the sample at up to 60 kHz. We obtained amino acid-type secondary structure information and used it to challenge a physically plausible atomistic model of PrP^Sc^ consisting of a 4-rung β solenoid recently proposed by us. In all cases, our model was compatible with the data. This study shows that elucidation of the structure of PrP^Sc^ is within reach using recombinant PrP^Sc^, NMR, and our model as a guiding tool.

## INTRODUCTION

The scrapie isoform of the prion protein (PrP^Sc^) was the first described and is the most notorious prion. PrP^Sc^ is an infectious protein that propagates by templating its conformation onto units of its alternatively folded conformer, the cellular prion protein (PrP^C^) (1-3). As it propagates through a mammalian brain, PrP^Sc^ causes fatal neurodegenerative diseases such as Creutzfeldt-Jakob disease (CJD) of humans, scrapie of sheep and goats, bovine spongiform encephalopathy (BSE) of cattle and chronic wasting disease (CWD) of cervids (1,2). The ability of PrP^Sc^ to propagate is encoded in its structure, and so are the strain and transmission barrier phenomena (1-3). We have recently proposed the first physically plausible atomistic model of PrP^Sc^ based on all the available experimental data, which consists of a 4-rung β solenoid (4RβS) and which is, in contrast to all previous models, stable through hundreds of nanoseconds of molecular dynamics simulations (4). The model offers a molecular explanation of PrP^Sc^ propagation as a process in which a section of the N-terminal flexible tail of PrP^C^ binds the C-terminal rung of PrP^Sc^, refolding to form three β-strands paired through hydrogen bonds with the three “unpaired” β-strands of the rung, which are ready set to form fresh hydrogen bonds and act as a template. Subsequent unfolding/refolding of the globular domain proceeds to form three additional β-strands to complete an additional rung, and so forth until completion of a new PrP^Sc^ subunit, whose C-terminal rung, with unpaired β-strands now becomes the new templating edge (3,4). Our model can also explain PrP^Sc^ strains, subtypes of PrP^Sc^ of a given sequence with distinct, stable biological and biochemical properties (1,2). PrP^Sc^ strains are believed to be PrP^Sc^ conformers featuring minor structural variations. Our model can accommodate such minor variations that can consist of subtle differences in the threading of β-strands and, more likely, the conformation of the loops connecting them (4). Our model also offers a simple solution to the steric conundrum of how the bulky side glycans jutting out from PrP^C^ pile up as it refolds into an amyloid, very difficult to account for with parallel-in-register (PIRIBS) models (5-7).

It is however obvious that despite these positive features and agreement with all the low-resolution structural constraints available (8-10), the model should be challenged against experimental structural constraints to validate it. These could come from solid state NMR spectroscopy (ssNMR). The advent of methods to prepare recPrP^Sc^ (11,12) has opened the possibility of generating the isotope-labelled material required for such studies. However, the relatively low output of the conversion methods used, all based on protein misfolding cyclic amplification (PMCA) techniques (13) has been a limitation, given that relatively large amounts (milligram quantities) of sample are required for ssNMR studies. We have recently developed a method, protein misfolding shaking amplification (PMSA), which circumvents this limitation as it allows facile production of milligram quantities of *bona fide*, infectious recombinant PrP^Sc^ (recPrP^Sc^). This in turn opens for the first time the possibility of applying ssNMR to the study of the structure of PrP^Sc^ (14). Given the complexity of ssNMR spectra of amyloids, typically featuring broad, overlapping signals, as a consequence of structural heterogeneity and peak crowding, solving the structure of recPrP^Sc^ will likely advance piecemeal and will require diverse isotopic labelling strategies. If our model proves to be able to accommodate the structural constraints derived from these studies as they begin to become available, it could be very useful as it might provide a working scaffold to integrate structural constraints, as opposed to having to construct models *ab initio* from likely limited ssNMR datasets. Here we describe a first study of uniformly labelled (^15^N,^13^C)-rec Bank Vole (BV) PrP(109I)^Sc^ treated with proteinase K (PK) whose results were used to challenge and refine our previously reported computational model.

## RESULTS

### Limited proteolysis of recBVPrP^Sc^(109I)

We used PMSA to generate unlabeled and uniformly labelled (U-^13^C,^15^N)-BVPrP^Sc^(109I)23-231. PMSA relies on shaking a PrP substrate in the presence of defined co-factors, zircone beads and a seed of preformed PrP^Sc^ (14). Our procedure includes a PK-treatment step at the end of the conversion process to ensure elimination of any non-converted PrP substrate. This material has been described in a previous publication (14) and is highly infectious, with attack rates of 100% and titers of 6.34·10^4^/μg of PrP in TgVole (1x). We carried out a full biochemical characterization of this material, to complete the partial one described in our previous study (14). We first analyzed the pattern of PK-resistant fragments using SDS-PAGE. Given that bands in the 5-10 kDa region are not well separated in the 4-12% commercial Tris/glycine SDS-PAGE gels that we used previously (14), we subjected the samples to 15% Tris/glycine gels, which allowed a much better separation and visualization of bands in this region, (Fig. 1A,B). With this system, we detected the known main fragments with apparent MWs of ∼15.5 and 9.5 kDa, respectively, but also fainter ones with MWs of ∼13, 11 kDa, and a faint, smeary band centered around ∼6 kDa (Fig. 1A). Next, we carried out a complete mass spectrometry-based analysis of the samples, which showed an excellent agreement with the SDS-PAGE-based analyses. PK-treated samples were sedimented by centrifugation, and the pellets denatured in 6 M Gdn/HCl and injected into a nanoHPLC coupled to an ESI-TOF detector. Fig. 1C shows the deconvoluted spectrum obtained, which allows an accurate identification of the PK-resistant fragments, with differences between experimental and theoretical mass values of +/– 1 Da. The main peaks correspond to fragments N_97_-S_231_, Q_98_-S_231_, and G_92_-S_231_ (the ∼15 kDa band seen in the SDS-PAGE gels); fragments N_153_-S_231_ and M_154_-S_231_ (the ∼9.5 kDa band seen in the gels); S_135_-S_231_ and F_141_-S_231_ (the ∼11 kDa band in the gels); N_97_-N_153_, N_98_-N_153_ and Q_98_-N_153_ (corresponding to the less intense ∼6 kDa band in the gels) and N_97_-F_141_ and N_97_-G_142_. Several additional small peaks could be also identified (Fig. 1C), and a few peaks remained unidentified. This fragmentation pattern agrees well with our model, although some adjustments of its original version, designed for brain-derived murine PrP^Sc^ (MoPrP^Sc^) (4) were necessary (Fig. 2). Thus, the predominant cleavages at N_97_/Q_98_ in recBVPrP^Sc^(109I) differ from the more prevalent cleavage at position G_90_ seen in mouse brain-derived RML type PrP^Sc^, which was used to build our model (4,8-10). Such a smaller main PK-resistant core, reminiscent of Drowsy-type PrP^Sc^ strains (15), has been described in recPrP^Sc^ species before (16). Also, while cleavages at S_135_, F_141_, G_142_, N_153_, M_154_, N_159_, and less conspicuously at A_117_, have similar counterparts in brain-derived PrP^Sc^ (4,8), we unexpectedly detected small peaks derived from cleavage before positions W_145_, E_146_, Y_149_ and R_151_. However, the relative abundance of the resulting fragments is extremely low.

**Figure 1.**
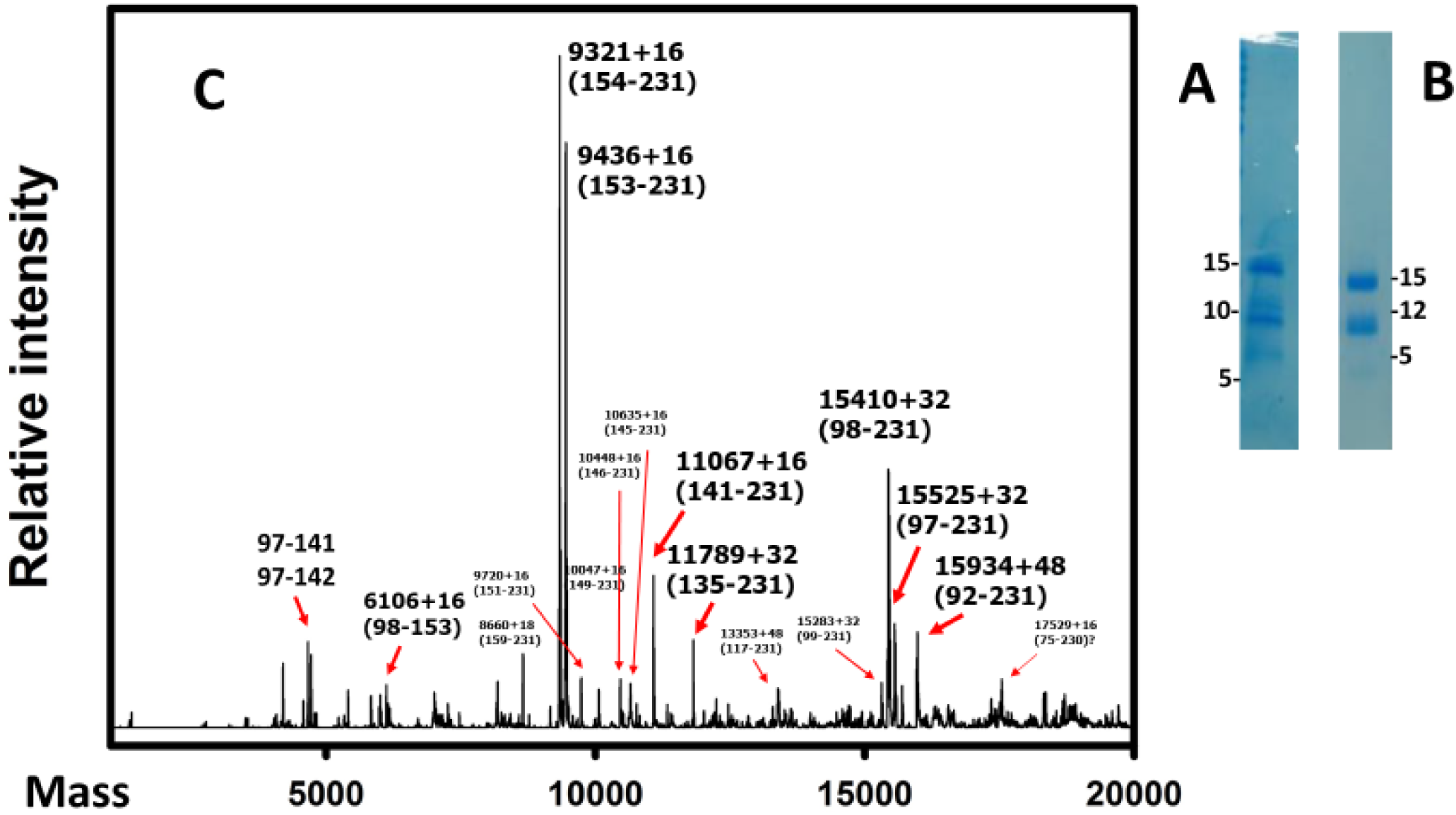
Fragmentation pattern of recBVPrP^Sc^ when subjected to limited proteolysis using PK. (**a**) Coomassie blue-stained SDS-PAGE performed under the conditions described in our previous study (14) or (**b**) using a 15% gel to improve separation of bands in the 5-10 kDa region. **(c)** Deconvoluted ESI-TOF spectrum; most peptides appeared as multiplets of M, M+16, M+32…corresponding to oxidation of methionine, likely occurring during PMSA. Samples were digested with 25 μg/ml of PK for 1 h at 37 °C in all cases.

**Figure 2.**
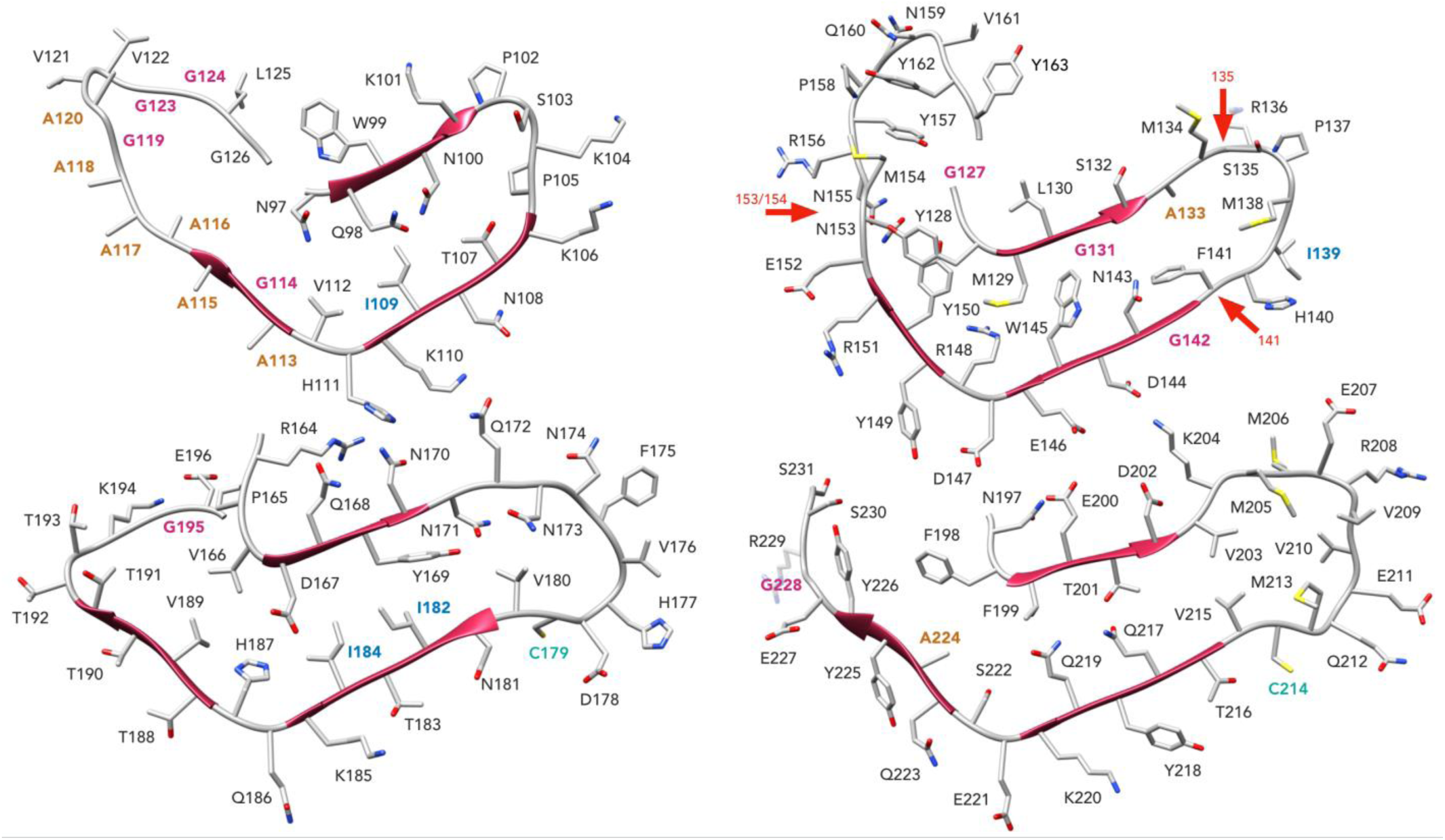
View of the model of the PK-resistant core of recBVPrP^Sc^(109I). Residues are displayed in each individual rung (1-4); the main PK cleavage sites identified by mass spectrometry are indicated with red arrows. Prolines are colored in purple; cystine is indicated in cyan; alanines are colored orange; isoleucines are colored blue; glycines are colored pink.

### 2D ^1^H-^13^C CP-HSQC ssNMR spectra of recBVPrP^Sc^

Next, we analyzed milligram quantities of uniformly labelled (U-^13^C,^15^N)-recBVPrP^Sc^ using ssNMR. We first obtained C-H CP HSQC spectra at ∼60 kHz MAS rate (Suppl. Fig. 1 and Fig. 3). Signals originating from the Hα–Cα atoms (see ref. 17 for the nomenclature used) of labelled amino acid residues appear within the region δC ≈ 47-65 ppm and δ_H_ ≈ 4-7 ppm (Fig. 3). We estimated the relative abundances of signals corresponding to residues that are located in β-strands as opposed to residues located in coil segments, based on their chemical shifts (18). It is clear from this spectrum that the majority of signals correspond to residues featuring β-sheet secondary structure. Choosing a value of 4.6 ppm in the ^1^H axis as the maximum for coil conformation (18), integration of the signals in the two resulting areas of the PK-treated (^13^C,^15^N)-BVPrP^Sc^ spectrum results in values of ∼85% β-sheet *vs*. ∼15% coil (Fig. 3). This distribution should be considered very cautiously, given that the frontier between both types of chemical shifts (18) is fuzzy; however, it is clear that the majority of signals detected derive from the β-stretches of the amyloid, with just a small fraction of signals arising from connecting loops with intermediate mobility, likely the shortest ones or positions within loops close to the rigid β-strands. The more mobile residues in the flexible loops become “invisible” to this NMR experiment, most likely as a consequence of fast molecular motion in the microscale, which renders those spins completely out of the window of rotational frequencies that match the Hartman-Hahn condition for the Cross Polarization (CP) period and therefore they do not generate signals in any spectrum based on CP (Suppl. Fig.2). It is also of note that the signals corresponding to β-strands showed a downfield spread, with signals reaching up to 7 ppm, something that to the best of our knowledge has not been reported. This might be a consequence of the high packing order and/or effects of the cross-beta CO–HN hydrogen bonds in the chemical shifts of the Hα/Cα resonances.

**Figure 3.**
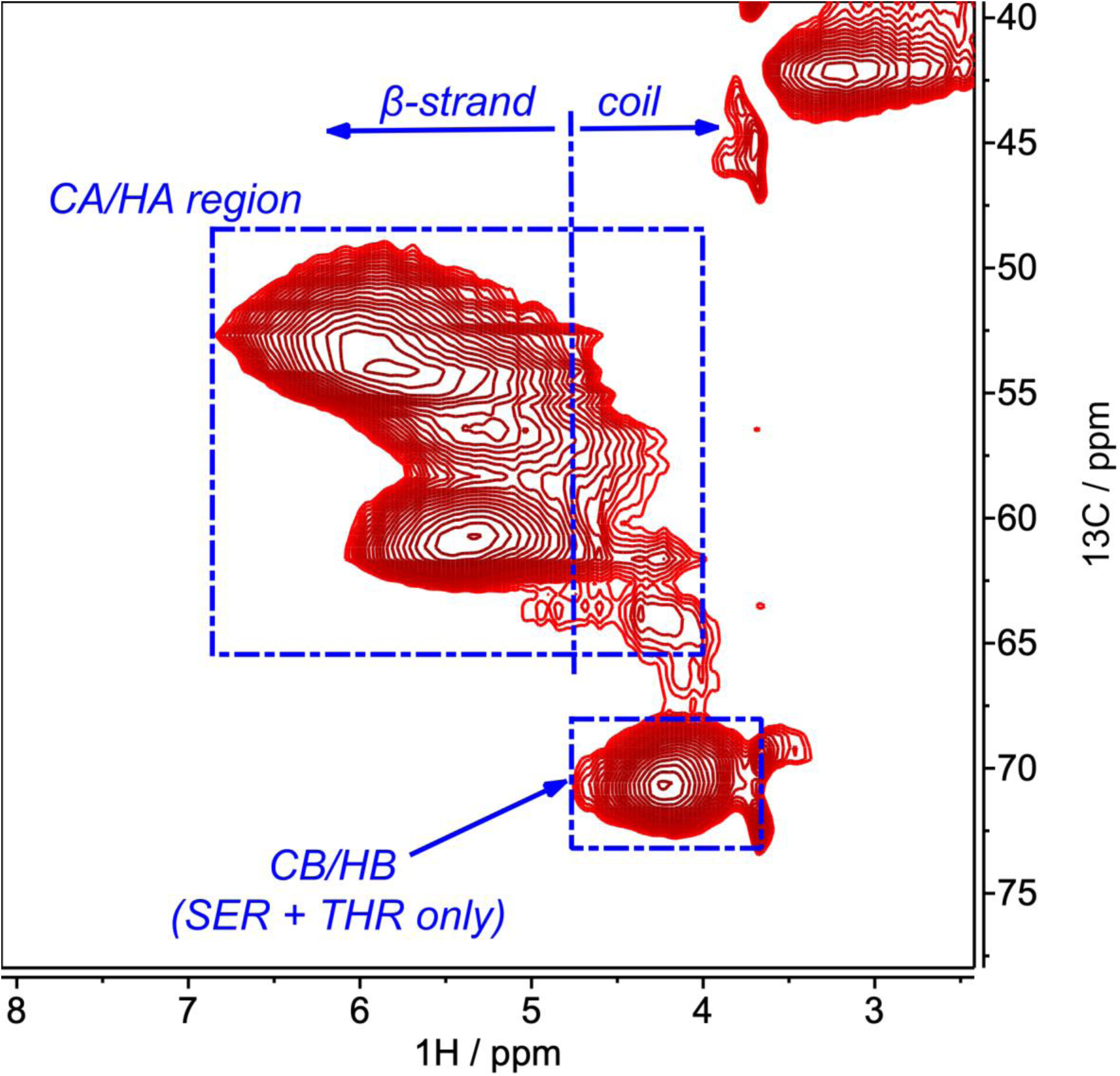
Hα/Cβ region of the 2D -^1^H-^13^C CP-HSQC spectra of PK-treated, uniformly labelled (U-^13^C,^15^N)-BVPrP^Sc^(109I).

### 2D ^13^C-^13^C ssNMR spectra of recBVPrP^Sc^

Next, 2D ^13^C-^13^C ssNMR measurements were performed on PK-treated (U-^13^C,^15^N)-BVPrP^Sc^(109I) samples. We performed both DREAM (Fig. 4) and DARR (Fig. 5) experiments to test for a variety of ^13^C-^13^C polarization transfer efficiencies that could enhance the detection of residues in a wide span of motional regimes. Fig. 4A shows a DREAM spectrum in which cross-peaks arise primarily from one-bond, intra-residue spin polarization transfers. Both ^13^C-^13^C experiments include a period of CP, and for the reason described in the previous section, the signals observed in these experiments should arise from residues in the most rigid parts of the protein fold (*e*.*g*. the β-sheets, *vide supra*), while residues that experience motions in the fast or intermediate time scale do not appear or have weak intensities (*e*.*g*. residues in the more flexible loops or in the terminal ends). In other words, the fact that an amino acid is observed in these ^13^C-^13^C spectra can be interpreted as an evidence suggesting that the residue is in a β-sheet or, at least, in a relatively rigid segment. In our sample signals are broad, as expected from an amyloid, and previously seen in similar spectra from PrP amyloid samples (19, 20). However, a wealth of relevant amino acid-class information can be extracted: **(i)** A minimum of four Ala peaks are observed in the 48-51/19-25 ppm chemical shift region of the DREAM spectrum, characteristic of β-strand secondary structure (18) (Fig 4C). No signals are observed for Ala residues with chemical shifts characteristic of coil or α-helical secondary structure. There are 8 Ala residues in PrP97-231, one in the C-terminal region (Ala_224_) and the other 7 in the N-terminal region: Ala_113_, Ala_115_, Ala_116_, Ala_117_, Ala_118_, Ala_120_ and Ala_133_ (Fig. 2). This means that at least 3 of these N-terminal Ala residues must feature a β-strand conformation, or, in other words, there are β-strands in the N-terminal region of BVPrP^Sc^(97-231), reaching a position as N-terminal as ∼Ala_116_. This has never been reported for any PrP amyloid, whose β-strands are confined to the ∼165-231 region (19,20), and is in excellent agreement with our model, in which A_113_ and A_115_ are in β-strand conformation, while A_116_ is at the edge of a β-strand and residues at the edge of β-strands display extended conformation over the course of MD simulations (Fig. 2). **(ii)** There are two sets of signals corresponding to Cβ/Cγ2correlations of Ile residues, in the 36-42/13-18 ppm region of the spectrum, featuring chemical shifts that are characteristic of β-strand and coil secondary structure, respectively (18) (Fig. 4B). Based on the relative intensities of the signals, and the fact that there are 4 Ile residues in BVPrP(97-231), it seems reasonable to conclude that 3 of these Ile residues are likely located in β-strands (δ_C_ ≈ 40/17 ppm) while 1 (δ_C_ ≈ 38.5/14 ppm) is located either in a short coil or at the edge of a long coil, near a β-strand. This agrees very well with our model (Fig. 2). **(iii)** Despite the fact that there are 13 Gly residues in BVPrP^Sc^(97-231), few CO/Cα signals corresponding to Gly residues are observed. These signals should appear around the characteristic 45/175 ppm region in the DARR spectrum (Fig. 5B) (18). The DARR spectrum obtained shows only ∼3-5 weak signals in this region. In general, Gly residues are conspicuously absent from β-strands (21). However, in our model 3 Gly residues are located in β-strands, and 2-3 more are in loops but in the vicinity of β-strands (Fig. 2). We recorded a second DREAM spectrum with 24 kHz MAS rate and 3 ms mixing time. Under these conditions, cross-peaks arising from Gly Cα/CO and Gly Cα/Cα correlations appear with opposite phase to the other cross peaks (Fig. 6) because of the particular sensitivity of DREAM to the experimental conditions (22). As seen in Fig. 6, we detected a clear group of signals corresponding to more than one Gly residue. Furthermore, in this spectrum we detected a few additional cross-peaks at ∼53/45 ppm that correspond to inter-residue correlations between Gly Cα and the Cα of contiguous residues. (**iv)** There is an intense signal corresponding to the Cα/Cβ correlation of several unresolved Thr residues at 61/70 ppm, with a weak neighboring signal corresponding to 1 Thr residue (Fig. 4A). Their chemical shifts values are compatible with both β-sheet and coil secondary structure. It is reasonable to conclude that the signals forming the intense unresolved peak come from a majority of Thr residues featuring β-sheet secondary structure. This agrees well with a vast majority, 8 of 11, Thr residues in BVPrP^Sc^97-231 being located in β-strands, and 3 in loops in close vicinity to β-strands (Fig. 2). **(v)** There are 3 clear, well resolved Ser Cα/Cβ cross-peaks in the 55-59/64-68 ppm region of the spectrum (Fig. 4D). Their distribution suggests that two of them might feature β-sheet conformation, while the other is likely more compatible with coil conformation (18), although this interpretation must be taken cautiously. Two additional well resolved peaks above these three are also compatible with the chemical shifts of Serine residues but might also correspond to Prolines (Fig. 4D). All of this is fully compatible with the model, which places 2 Ser residues in β-strands, and 1 additional Ser and two Pro residues at the very short Pro_102_-Pro_105_ connecting loop, likely to have little motion and thus be detectable. (Fig. 2). **(vi)** Two regions of the DARR spectrum display cross-peaks corresponding to aromatic amino acid residues: region 115-163 ppm for C-aromatic/C-aromatic correlations and region 30-60/115-138 ppm for correlations of the aliphatic Cα and Cβ carbons with the aromatic carbons. Signals corresponding to ∼6-8 Tyr residues could be unequivocally discerned from the diagonal peaks of Tyr carbons Cε (δ_C_ ∼117 ppm) and Cζ (δC ∼157 ppm), as these chemical shift values are privative of Tyr. Other peak groups in the aromatic subspectrum confirm this assignment, like those correlations between the Cα/Cε and the Cε/Cζ carbons (Fig. 5C). This agrees well with 6 Tyr residues located in β-strands in our model. Besides, a small number of additional, non-Tyr aromatic amino acid residues (Phe, Trp or His) can be discerned at 125-140 ppm (Fig. 5C), also in agreement with the presence of 2 Trp and 1 His in β-strands in the model (Fig. 2). **(vii)** Also, in this DARR spectrum, 2 sets of signals unequivocally corresponding to Cδ/Cζ (∼45/160 ppm) and Cγ/Cζ (∼28/160 ppm) correlations of Arg are seen (Fig. 5A). Each set includes ∼2 overlapping signals (Fig. 5A). In our model (Fig. 2), 2 of a total of 7 Arg residues are located in β-strands.

**Figure 4.**
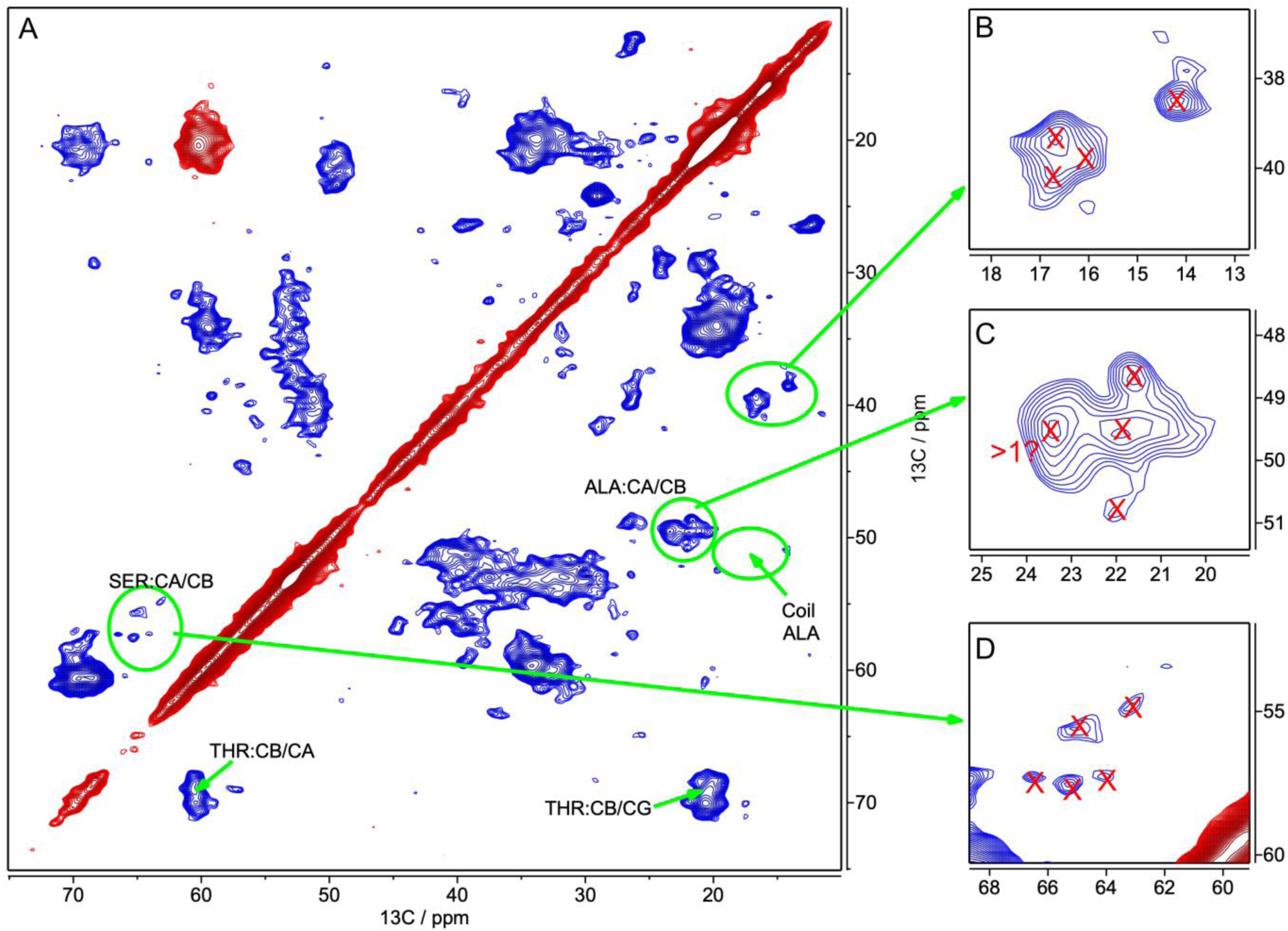
2D ^13^C-^13^C DREAM spectrum (45 kHz MAS, 3 ms mixing time) of PK-treated, uniformly labelled (U-^13^C,^15^N)-BVPrP^Sc^(109I). **(A)** Expansion of the aliphatic region (Cα, Cβ, Cγ…). **(B)** Expanded 36-42/13-18 ppm region, where Ile Cβ/Cγ2 correlation cross-peaks appear. **(C)** Expanded 48-51/19-25 ppm region, where Ala Cα/Cβ correlation cross-peaks appear. **(D)** Expanded 52-60/59-68 ppm region, where Ser Cα/Cβ correlation cross-peaks appear.

**Figure 5.**
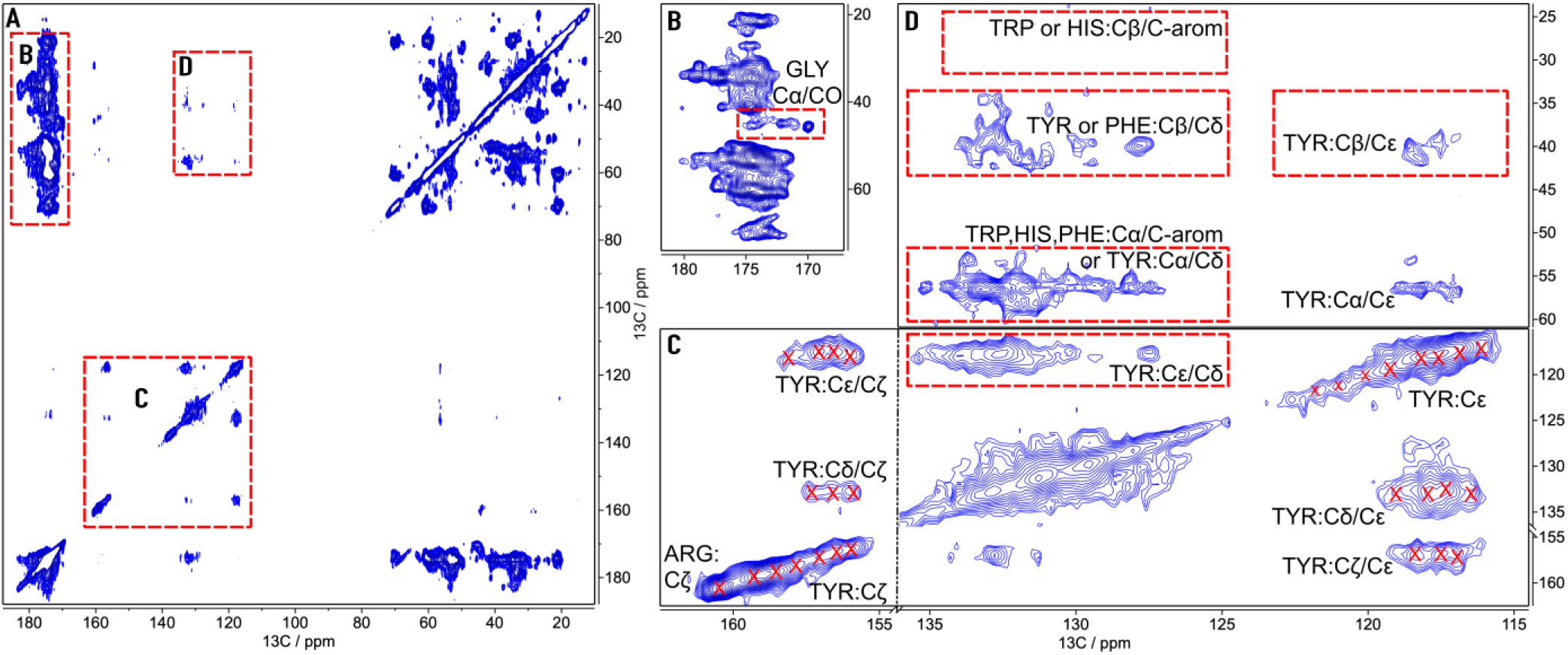
2D ^13^C-^13^C DARR spectrum of PK-treated, uniformly labelled (U-^13^C,^15^N)-BVPrP(109I)^Sc^, recorded at 20 kHz MAS rate with 80 ms mixing time. (A) Full spectrum: red rectangles contain the expanded regions. (B) Expansion of the C-aliphatic/CO region. Red rectangle contains the GLY:CA/CO correlations (≈3-5 peaks). (C) Expansion of the C-aromatic/C-aromatic region. A minimum of 6 TYR residues can be deduced from diagonal peaks and correlations of their Cδ, Cε and Cζ atoms. It is unclear if diagonal peaks in the range δC ∼ 120-123 ppm are additional TYR:Cε resonances or if they correspond to aromatic carbons from TRP and HIS residues. Contribution of other aromatic residues (PHE, HIS, TRP) to peaks in the region 125-130 ppm can not be discarded, nor confirmed, due to overlap of their typical chemical shift ranges (18). (D) Expansion of the C-aliphatic/C-aromatic correlations. Cross-peaks in the δC ∼ 115-120 ppm region (Cε) can be unequivocally assigned to TYR:Cα/Cε (δCα ∼ 57 ppm) and TYR:Cβ/Cε (δCβ ∼ 41 ppm).

**Figure 6.**
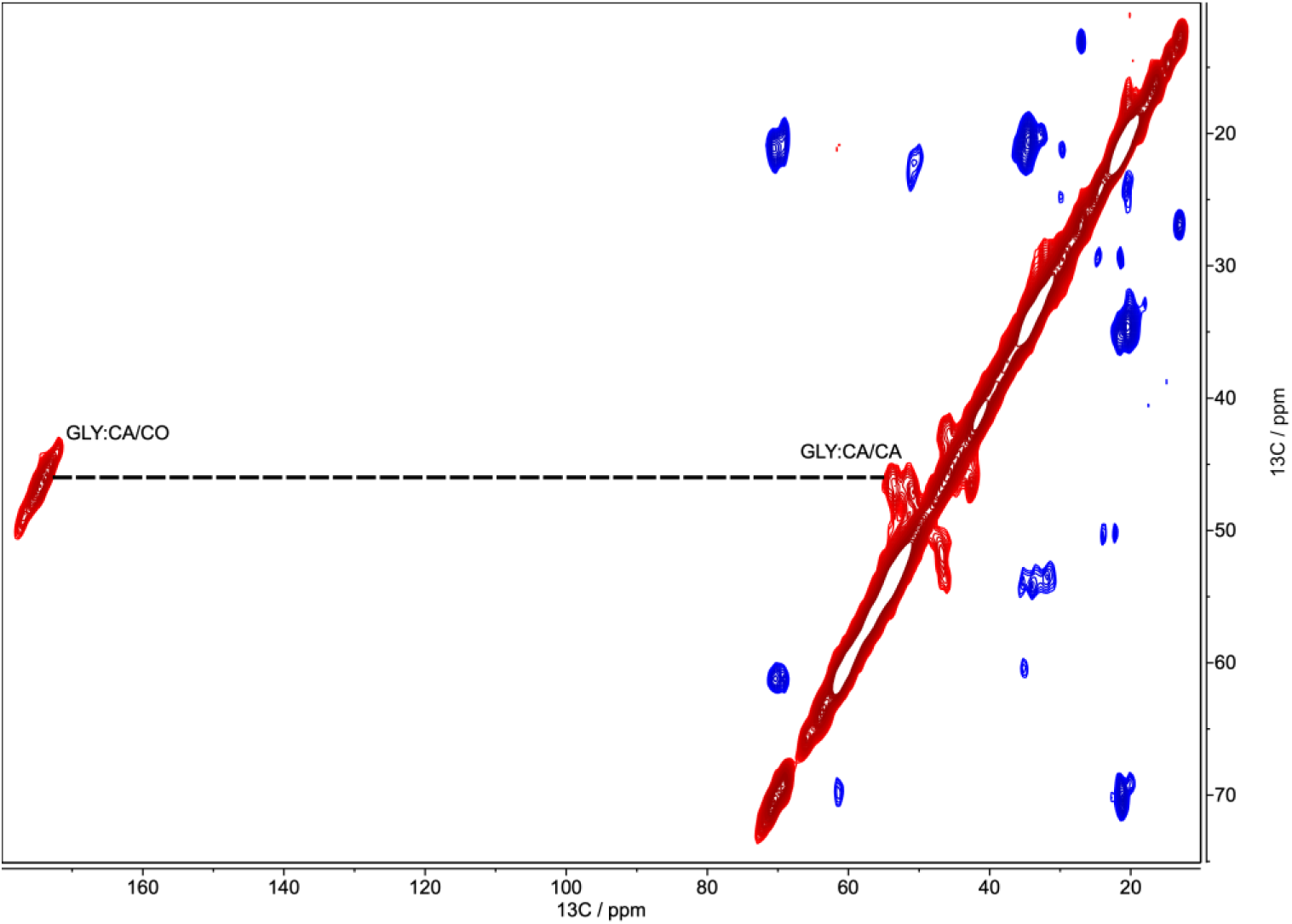
2D ^13^C-^13^C DREAM spectrum (24 kHz MAS, 3 ms mixing time) of PK-treated, uniformly labelled (U-^13^C,^15^N)-BVPrP^Sc^(109I).

### 2D ^1^H-^13^C and ^1^H-^15^N INEPT-HSQC spectra of recBVPrP^Sc^

A uniformly labelled PK-treated (U-^13^C,^15^N)-BVPrP^Sc^ sample was also subjected to ssNMR analysis with methods based on INEPT magnetization transfer and rotation at high MAS rates (2D ^1^H-^13^C and ^1^H-^15^N INEPT-HSQC experiments). Differently to what occurs with experiments based in CP, INEPT based experiments are more suitable to detect the resonances of the most mobile or flexible stretches or domains of the solid sample (Suppl. Fig. 2). Under these conditions, a small number of signals were detected in spectra of (U-^13^C,^15^N)-BVPrP^Sc^ when the sample was rotated at ∼60 kHz MAS rate. We identified some of the peaks in the 2D ^1^H-^13^C INEPT-HSQC spectrum as originating from natural abundance ^13^C of dextran and Triton X-100 molecules that are tightly bound to the recBVPrP^Sc^ (data not shown). Both compounds are used as co-factors during preparation of recBVPrP^Sc^ (see Online methods). We therefore focused our attention on the ^1^H-^15^N INEPT-HSQC spectra (Fig. 7), as all their signals unequivocally derive from amino acid residues in recBVPrP^Sc^ because neither Triton nor dextran have nitrogen atoms. Based on the distinctive chemical shift values of the side-chain H-N atoms, we can identify peaks corresponding to 2 Gln/Asn residues (δ_N_ ∼110-112 ppm and δ_H_ ∼ 7.1 and 7.8 ppm) and 1 Arg (δ_Nε_ ∼83 ppm and δ_Hε_ ∼ 7.4 ppm). Furthermore, cross-peaks in the 117-120 ppm/8.1-8.8 ppm area are compatible with backbone HN of 2-3 Ser and/or Thr residues. A group of a minimum of 4-5 additional unresolved signals is also present in the δ_N_ ∼119-127 ppm/ δ_H_ ∼ 8.0-8.5 ppm region (Fig. 7). Finally, an intense signal centered at 129.5 ppm/7.8 ppm, a chemical shift value typical of N atoms in C-terminal amino acid residues was also seen. The intensity of these INEPT-HSQC signals decreased very substantially as the rotation MAS rate was decreased, virtually disappearing below 20 kHz (data not shown). This dependence on the MAS rate is also in agreement with those residues being in highly flexible segments of the molecule (Suppl. Fig. 2).

**Figure 7.**
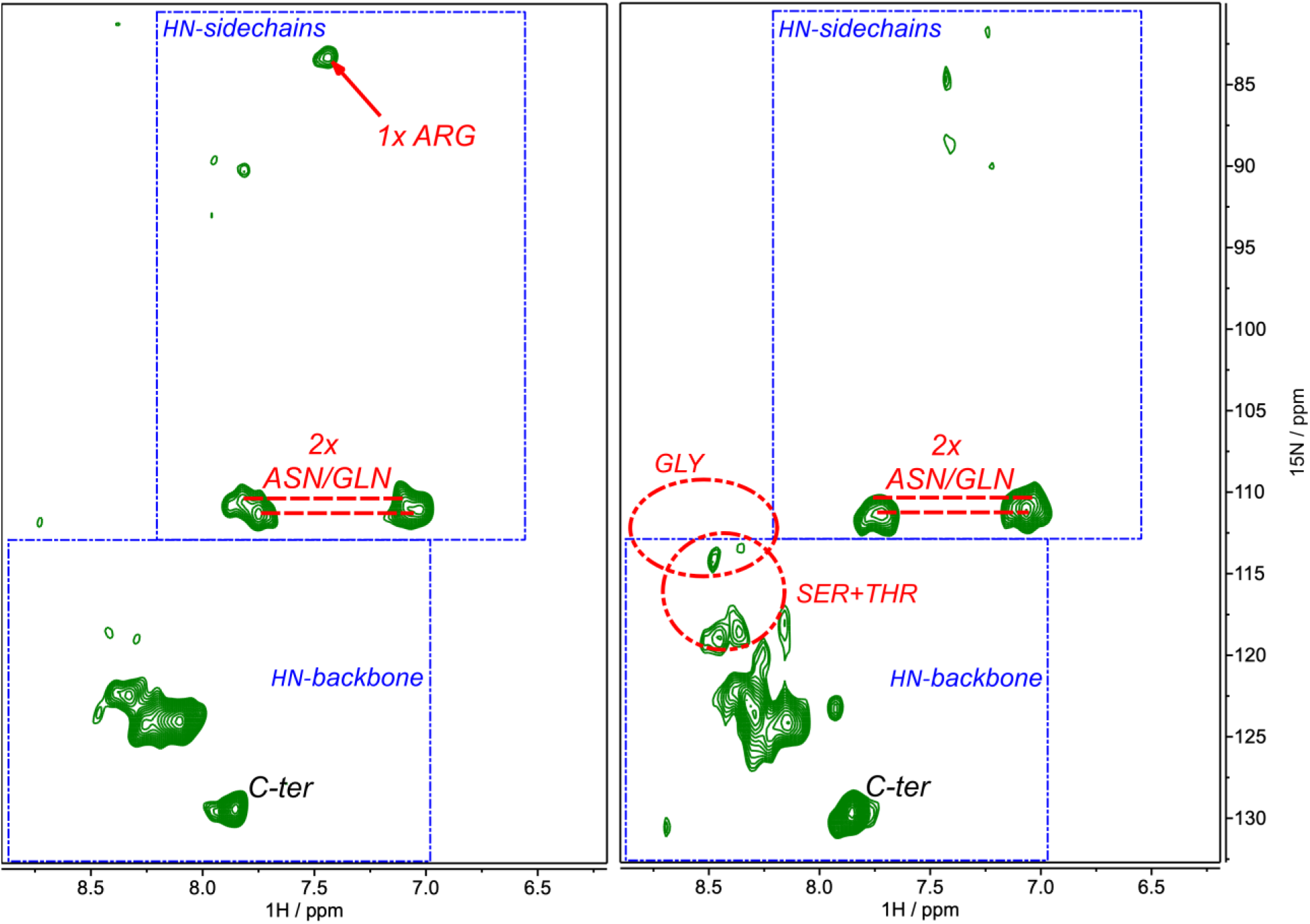
Sections of 2D ^1^H-^15^N INEPT-HSQC spectra of PK-treated, uniformly labelled (U-^13^C,^15^N)-BVPrP^Sc^(109I) obtained at a MAS rate of **(a)** 45 kHz, and **(b)** 60 kHz.

### Model building and Molecular dynamics simulations

We adapted our 4RβS full atomistic model of MoPrP^Sc^ (4) to recBVPrP^Sc^(109I). For that purpose, we adapted the threading scheme of murine PrP^Sc^ to the PrP sequence of bank vole 109I. Single residue variations were switched from the mouse sequence to the bank vole sequence. Only a single major rearrangement was necessary, involving the N-terminal rung, which is shorter after PK cleavage in bank vole (Fig. 2). The 3D structure of recBVPrP^Sc^(109I) was then built by following the protocol reported in using the adapted threading scheme (see Online methods for details). The resulting structure agrees well with the PK cleavage pattern of recBVPrP^Sc^(109I) (Fig. 1) except for minor nicks affecting predicted β-strands Gly_142_-Glu_146_ and Tyr_149_-Arg_151_.

Next, a model of the tetramer was built by stacking four recBVPrP^Sc^(109I) subunits vertically and was subsequently subjected to Molecular Dynamics Simulation. The percentage of β-sheet along the course of the 100 ns simulation (mean of N = 3) changed from an initial value of 39 ± 1 % (mean of the first 10 ns, ± standard deviation) to a final value of 37 ± 1 % (mean of the last 10 ns, ± standard deviation). This minor change is indicative of high stability, similar to that of our mouse 4RβS tetramer model (4).

## DISCUSSION

Availability of milligram quantities of *bona fide* recPrP^Sc^ opens the possibility of using a variety of biophysical techniques, in particular ssNMR, to the analysis of the structure of this quintessential prion. In fact, full elucidation of the structure of recPrP^Sc^ is a plausible goal for the first time. Here, we show a first analysis of uniformly labeled PK-treated recPrP^Sc^. We have performed a variety of ssNMR experiments at moderate (20-24 kHz) and higher (50-60 kHz) MAS rates, which allowed us to acquire 2D spectra involving not only ^15^N and ^13^C but also ^1^H nuclei. These spectra display broad signals, as seen for other amyloids, given that they all have in common some degree of conformational diversity within a given sample (19,20,23). The exception to this is the functional Het-s prion domain, whose conformation has been subjected to natural selection and is therefore very well defined, which results in ssNMR spectra with sharp peaks (24). However, clearly this is not the case for PrP^Sc^, as it is an amyloid conformer with no expected function, and without the stringent selective pressures associated to one. Nevertheless, in spite of the severe signal overlap, we have gleaned a substantial amount of unprecedented structural information from the ssNMR spectra. We have used this information to challenge our full atomistic PrP^Sc^ model (4) and found it to be fully compatible with these new high-resolution pieces of data. As an example, one important conclusion derived from the DARR spectra is that β-strands extend into the N-terminus of the recPrP^Sc^ PK-resistant core, as at least two of the Alanines in the Ala_113_-Ala_120_ sequential stretch are in β-sheet conformation (Fig. 4). This agrees with prior hydrogen/deuterium exchange studies coupled to fragmentation and mass spectrometry that have shown slow exchange rates typical of β-strands in overlapping pepsin fragments covering most of the ∼90-225 sequence of both brain-derived and recombinant PrP^Sc^ samples (25,26). Now, direct spectroscopic evidence of these β-strands is presented. Of note, our model features three Ala residues in a β-strand spanning Val_112_-Ala-Gly-Ala-Ala_116_. This β-strand looks *a priori* unlikely, given that it contains 4 residues with very low propensity to participate in β-strands (21), of a total of 5 residues. In contrast, DARR spectra of conventional murine and ovine PrP amyloids obtained by Tycko *et al*. and Müller *et al*. (19,20) showed two distinct sets of Ala Cα/Cβ signals, corresponding to residues featuring both β-sheet and non-β-sheet (loop or α-helical secondary structure), in agreement with the fact that in these amyloids the majority of the Ala residues are outside of the β-sheet-rich core, which spans from position ∼155 to the C-terminus. The fact that the experimental data presented here support the existence of this theoretically unlikely β-strand militates in favor of our model. In contrast, our spectroscopic data suggest that at least one Ile residue is not located in a β-strand (Fig. 4), despite the fact that Ile has the second highest propensity to participate in β-strands (21). And this agrees with our model, which locates Ile_139_ in a connecting loop.

Analysis under conditions more suitable for soluble state samples or highly flexible semi-solid samples, *i*.*e*., 2D ^1^H-^15^N INEPT-HSQC experiments recorded at high MAS rates, showed that a few amino acid residues of PK-treated recBVPrP^Sc^ are located in regions with very high mobility. While sequential information could not be obtained, we were able to identify a few residues based on their distinct chemical shift values. These include 2 Gln and/or Asn residues, 1 Arg and 2-3 Ser and/or Thr residues among a total of ∼10-12 residues. Smirnovas *et al*. and Noble *et al*. have reported deuterium/hydrogen exchange studies suggesting that a short C-terminal (but not N-terminal) sequential stretch is substantially flexible, with high exchange rates (25,26). The C-terminal stretch of BVPrP is: Ser_231_-Ser-Arg-Gly-Glu-Tyr-Tyr_225_. Our ^1^H-^15^N INEPT-HSQC spectra contained unequivocal signals corresponding to a minimum of 2 Serines and one Arginine. A possible weak Glycine peak was also seen (Fig. 7). A sizeable fraction of our PK-treated recBVPrP^Sc^ sample has a nick at positions Asp_153_ and Met_154_. Our model predicts a loop spanning Asp_153_ to Pro_165_, based on the conspicuous accessibility of this region to PK (Fig. 1) and the presence of the β-strand breaking amino acid residues Pro_158_ and Pro_165_. A nick at Asp_153_ and/or Met_154_ might disrupt such loop, allowing increased motion of its C-terminus: Asp_154_-Arg-Tyr-Pro-Asp-Glu-Val_160_. The combination of amino acid residues in these two stretches agrees very well with the signals seen in our 2D ^1^H-^15^N INEPT-HSQC spectrum.

The data we have been able to extract from ssNMR spectra do not allow discarding a parallel-in-register (PIRIBS) architecture, that has been alternatively proposed for PrP^Sc^ (5). A PIRIBS structure in which several Ala residues within the A_114_-A_121_ region are part of a β-strand, one of the 4 Ile residues within the Q_97_-S_231_ main PK-resistant core is located within a loop, and the majority, but not all Gly residues are excluded from β-strands, is theoretically possible, although the currently available PIRIBS model of PrP^Sc^ would have to be re-threaded, as in its current version it does not fully comply with the data presented here.

In conclusion, we have carried out a first set of ssNMR analyses of a *bona fide* recombinant PrP^Sc^ prion. The structural constraints obtained are in excellent agreement with our 4-rung β-solenoid model of PrP^Sc^. We will continue to use ssNMR to obtain additional structural constraints and will continue to use these to challenge and refine our model. A final, definitive elucidation of the structure of PrP^Sc^ is thus, for the first time, near.

## METHODS

### Production of recombinant protein

RecBVPrP(109I)23-231 was expressed by competent E. coli Rosetta (DE3) bacteria harboring pOPIN E expression vector containing the wild type I109 bank vole *Prnp* gene (https://pubmed.ncbi.nlm.nih.gov/29094186). Bacteria from a glycerolate maintained at –80 °C were grown in a 250 ml Erlenmeyer flask containing 50 ml of LB broth overnight. The culture was then transferred to two 2 L Erlenmeyer flasks containing each 500 ml of minimal medium supplemented with 3 g/L glucose, 1 g/L NH_4_Cl, 1 M MgSO_4_ (1 ml/L), 0.1 M CaCl_2_ (1 ml/L), 10 mg/ml thiamine (1 ml/L) and 10 mg/ml biotin (1 ml/L). For production of uniformly labelled (U-^13^C,^15^N)-PrP, glucose and NH_4_Cl were substituted by (^13^C)-glucose and ^15^NH_4_Cl (Cortecnet, Paris) as the sole carbon and nitrogen sources. When the culture reached an OD_600_ of ∼0.9-1.2 AU, Isopropyl β-D-1-thiogalactopyranoside (IPTG) was added to induce expression of PrP overnight under the same temperature and agitation conditions. Bacteria were then pelleted, lysed, inclusion bodies collected by centrifugation, and solubilized in 20 mM Tris-HCl, 0.5 M NaCl, 6 M Gdn/HCl, pH = 8. Although the protein does not contain a His-tag, purification of the protein was performed with a histidine affinity column (HisTrap FF crude 5 ml, GE Healthcare Amersham) taking advantage of the natural His present in the octapeptide repeat region of PrP. After elution with buffer containing 20 mM Tris-HCl, 0.5 M NaCl, 500 mM imidazole and 2 M Gdn/HCl, pH = 8, the quality and purity of protein batches was assessed by BlueSafe (NZYTech, Lisbon) staining after electrophoresis in SDS-PAGE gels. Finally, Gdn/HCl was added, to a final concentration of 6 M, for long-term storage of purified protein preparations at –80 °C.

### Conversion of recBVPrP(109I)23-231 to recBVPrP(109I)^Sc^

Conversion was carried out by PMSA as previously described (14). Briefly, the purified rec-PrP stored in buffer containing 6 M Gdn/HCl (*vide supra*) was diluted 1:5 in phosphate buffered saline (PBS) and dialyzed against PBS at 1:2,000 ratio for 1 h at room temperature. The dialyzed sample was centrifuged at 19,000 g for 15 min at 4 °C and the supernatant used for substrate preparation. Rec-PrP concentration in the supernatant was measured (BCA protein assay kit, Thermo Scientific) and adjusted to the working concentration, which, unless otherwise indicated, was of 20 μM. The protein was then mixed at a 1:9 ratio with conversion buffer (150 mM NaCl, 10 g/ L Triton X-100, and 0.5% w/v of dextran sulfate sodium salt, from *Leuconostoc spp*. with sizes ranging from 6,500 to 10,000, Sigma-Aldrich in PBS). The substrate was aliquoted and stored at –80 °C until required. PMSA was performed by transferring 20 ml of the rec-PrP substrate to a 50 ml Falcon tube containing 2.8 g of washed 1 mm zirconia/silica beads (11079110Z, BioSpec Products Inc.). The tube was placed in a Thermomixer (Eppendorf) and incubated at 39 °C and 7,000 rpm in a continuous mode for 24 h.

### Proteinase K digestion

Samples were digested by addition of PK (Roche) from a concentrated stock solution to a final concentration of 25 μg/ml and incubation at 42 °C for 1 h. The sample was then immediately centrifuged at 19,000 g at 4 °C for 30 min, the supernatant was discarded and the pellet resuspended and washed with 1 ml of PBS. After a further 30 min at 19,000 g at 4°C, the supernatant was discarded.

### Analysis of PK-induced fragmentation

PK-treated recBVPrP^Sc^(109I) pellets were dissolved in Laemmli buffer and heated at 95 °C for 10 min, followed by SDS-PAGE in home-made 15% Tris/glycine gels or commercial 4-12% Tris/glycine gels (NuPage, Thermo-Fisher). After electrophoresis, gels were stained with BlueSafe Coomassie stain (NZYTech, Lisbon). For mass spectrometry-based analysis, the pellets were resuspended in 50 μl of 6M Gdn/HCl with 3 pulses (5 s each) of a tip sonicator and incubated at 37 °C for 1 h. TFA was added to a final concentration of 1%. Samples (4 μl) were injected to a micro liquid chromatography system (Eksigent Technologies nanoLC 400, SCIEX) coupled to a high speed Triple TOF 6600 mass spectrometer (SCIEX) with a micro flow source, and equipped with a silica-based reversed phase column Chrom XP C18 150 × 0.30 mm, 3 mm particle size and 120 Å pore size (Eksigent, SCIEX). A YMC-TRIART C18 trap column was connected prior to the separating column, on-line (3 mm particle size and 120 Å pore size, YMC Technologies, Teknokroma). After sample loading at and washing with 0.1% formic acid in water to remove Gdn/HCl and other non-peptide components of the sample, the flow was switched on to the analytical column and separation proceeded at a flow rate of 5 µl/min with a solvent system consisting of 0.1% formic acid in water as mobile phase A, and 0.1% formic acid in acetonitrile as mobile phase B. Peptides were separated over 40 min with a gradient ranging from 2% to 90% of mobile phase B. Data acquisition was performed in a TripleTOF 6600 System (SCIEX, Foster City, CA) using a Data dependent workflow. Source and interface conditions were the following: ionspray voltage floating (ISVF) 5500 V, curtain gas (CUR) 25, collision energy (CE) 10 and ion source gas 1 (GS1) 25. Instrument was operated with Analyst TF 1.7.1 software (SCIEX, USA). Switching criteria was set to ions greater than mass to charge ratio (m/z) 350 and smaller than m/z 1400 with charge state of 2–5, mass tolerance 250 ppm and an abundance threshold of more than 200 counts (cps). Former target ions were excluded for 15 s. The instrument was automatically calibrated every 4 hours using tryptic peptides from PepCalMix as external calibrant. For data analysis, the sample TIC was opened using the Peak View 2.2 software that allows protein reconstruction. The LC-MS Peptide Reconstruct feature uses a peak finding algorithm to identify groups of peaks that form isotope series and charge series. Protein deconvolution was carried out between 800 to 20000 Da.

### Solid State NMR measurements (ssNMR)

*Sample preparation:* (U-^13^C,^15^N)-recBVPrP^Sc^(109I) solution, as obtained by PMSA was treated with 25 μg/ml of PK (*vide supra*). The sample was then centrifuged at 9,000 *g* for 1 h at 4 °C in 85 ml OAK polycarbonate (Nalgene) tubes using a FiberLite F15-6×100 rotor (Piramoon Technologies, Inc.) in a Beckman Sorvall Legend XTR/230V ultracentrifuge. The supernatant was carefully removed, and the pellet washed with milliQ water, centrifuged under the same conditions for additional 15 min and supernatant removed to obtain the final pellet. The sample was then prepared in a 1.3 mm rotor for ssNMR measurement using several tools included in a Bruker solid sample preparation kit. The final pellet was resuspended in 50 μl of water by repeated pipetting and transferred to the loading funnel with the 1.3 mm rotor inserted. This assembly was placed in a home-built desiccation chamber containing CaCl_2_ until all the water evaporated, leaving behind thin PrP^Sc^ scales around the border of the 1.3 mm rotor. Using the Bruker loading rod, the scales were carefully introduced and compacted into the rotor, together with 2 μl of milliQ water to obtain a hydrated sample. The rotor was capped and placed in the solid NMR probe. Solid NMR experiments were measured at 278 K in a Bruker NEO 17.6 T spectrometer (proton resonance 750 MHz) spectrometer equipped with a ^1^H/^13^C/^15^N triple resonance solid probe for 1.3 mm zirconia rotors and an available range of MAS rates from 8 to 67 kHz. The spectrometer control software was TopSpin 4.0. All the spectra were processed with MestreNova software v14.0 (Mestrelab Research Inc.). Carbon-13 chemical shifts were referenced to the CA signal of solid glycine at 43.5 ppm. Nitrogen-15 chemical shifts were referenced to the ^15^N peak of a solid ^15^N labelled sample of glycine at 35.0 ppm. Proton chemical shifts were referenced to the intense water peak at 4.7 ppm. The following experiments were carried out:

2D ^1^H-^13^C CP-HSQC spectra were measured at 50 kHz MAS (pulse sequence *hCH2D*.*dcp* of the Bruker library). The ^1^H and ^13^C carriers were centered at 2.27 and 80 ppm, respectively. The spectral width in ^1^H and ^13^C dimensions were 19.6 and 175 ppm, respectively. The initial proton/carbon cross polarization contact time was 0.7 ms and the proton contact pulse power used an ascending linear ramp of 50%. The final carbon/proton cross polarization contact time was 0.4 ms and the proton contact pulse power used a descending linear ramp of 50%. MISSISSIPPI water suppression pulses were applied in proton at 15 kHz during 7.66 ms. The number of points acquired in t_2_ and t_1_ dimensions were 588 and 300, respectively. The FID was acquired under heteronuclear decoupling at 20 kHz with WALTZ-16 for both ^13^C and ^15^N nuclei. The inter-scan delay was 1.4 s and the number of scans was 16.

2D ^1^H-^15^N INEPT-HSQC spectra were measured at 50 kHz MAS (pulse sequenc*e hNH2D*.*hsqc* of the Bruker library). The ^1^H and ^13^C carriers were centered at 6 and 119 ppm, respectively. The ^1^H and ^13^C spectral widths were 19.81 and 43.86 ppm, respectively. The INEPT delays were optimized for relaxation losses using a nominal value of scalar coupling ^1^J_NH_ of 100 Hz slightly over the expected value of 95 Hz. MISSISSIPPI water suppression pulses were applied in proton at 15 kHz during 1.66 ms. The number of points acquired in t_2_ and t_1_ dimensions were 588 and 128, respectively. The FID was acquired under ^15^N heteronuclear decoupling at 20 kHz with WALTZ-64. The inter-scan delay was 1 s and the number of scans was 80.

2D ^1^H-^13^C INEPT-HSQC spectra were measured at 45 or 60 (pulse sequenc*e hCH2D*.*hsqc* of the Bruker library). The ^1^H and ^13^C carriers were centered at 2.27 and 75 ppm, respectively. The ^1^H and ^13^C spectral widths were 19.6 and 180 ppm, respectively. The INEPT delays were optimized for relaxation losses using a nominal value of scalar coupling ^1^J_CH_ of 200 Hz, a value that is over the expected value of 145 Hz. MISSISSIPPI water suppression pulses were applied in proton at 15 kHz during 1.66 ms. The number of points acquired in t_2_ and t_1_ dimensions were 323 and 220, respectively. The FID was acquired under ^13^C and ^15^N heteronuclear decoupling at 20 kHz with WALTZ-16. The inter-scan delay was 1.6 s and the number of scans was 48.

2D ^13^C-^13^C DARR spectra were measured at 20 kHz MAS (pulse sequence *cpSPINDIFF* of the Bruker library). The ^13^C carrier was centered at 110 ppm and the spectrum covered a spectral width of 220 ppm in each carbon dimension. The ^1^H carrier was centered over the water at 4.7 ppm. The initial proton/carbon cross polarization contact time was 2 ms and the proton contact pulse power used an ascending linear ramp of 50%. During the mixing time of the sequence a DARR ^1^H pulse was applied at 20 kHz. The number of points acquired in t_2_ and t_1_ dimensions were 1666 and 300, respectively. The FID was acquired under heteronuclear proton decoupling at 110 kHz with SPINAL-64. The inter-scan delay was 2.5 s and the number of scans was 88.

2D ^13^C-^13^C DREAM spectra were measured at 45 and 24 kHz MAS (pulse seque*nce hCC_dream2D-cp* of the Bruker library). The ^13^C carrier was centered at 110 ppm and the spectrum covered a spectral width of 220 ppm in each carbon dimension. The ^1^H and ^15^N carriers were centered at 3.5 and 120 ppm, respectively. The DREAM mixing time was 3 ms and consisted in a ^13^C adiabatic pulse of frequency given by 0.45 x w_r_, which in our spectra correspond to a frequency of 20.3 (/10.8) kHz at 45 (/24) kHz MAS. The ^13^C adiabatic pulse was applied at a frequency of 50 ppm and simultaneously with ^1^H CW decoupling at 110 kHz. The number of points acquired in t_2_ and t_1_ dimensions were 1000 and 600, respectively. The FID was acquired under heteronuclear decoupling in ^1^H at 110 kHz with SPINAL-64 and in ^15^N at 20 kHz with WALTZ-16. The inter-scan delay was 2 s and the number of scans was 32.

### Model building and simulations

The 4RβS model of bank vole (*Myodes glareolus*) PrP^Sc^ was constructed by adapting the threading scheme of *Mus musculus* PrP^Sc^ reported in (4) to the PrP sequence of bank vole M109I. Mouse PrP and bank vole PrP share 96% sequence identity computed from residue 97 to 230 (BV numbering). In the employed scheme, single residue variations were switched from the mouse sequence to the bank vole sequence; only a single major rearrangement was necessary, that involves the N-terminal rung, that is shorter after PK cleavage in bank vole. The 3D structure of BVPrP^Sc^ was built by following the protocol reported in (4) using the adapted threading scheme. Briefly, the 4 rung scaffold was obtained from *Dickeya dadantii* pectate lyase (PDB accession code: 1AIR); the core residues were replaced with BVPrP ones using UCSF Chimera (27) and original loops were replaced with bank vole loops generated using MODELLER (28). Protein topology was generated using Gromacs 2018 with Amber 99SB-ILDN force field (30) and the structure was energy minimized in vacuum using the steepest descent algorithm. The output conformer was placed in a cubic box, solvated with spc216 water molecule (TIP3P), neutralized with 8 Cl^−^ ions and energy minimized again. This structure was then refined by correcting the Ramachandran outliers and the low probability rotamers using Coot (31) and Chimera. Absence of steric clashes was confirmed using the specific Clashes/Contacts detection tool in Chimera: VDW-overlap threshold was set at 0.6 Å, subtracting 0.4 Å to account H-Bonds and ignoring contacts of atoms less than 4 bonds apart. Tetramer structure was obtained by stacking four monomers in a head-to-tail fashion. PrP^Sc^ tetramer topology was generated in Gromacs 2018 with Amber99SB-ILDN force field. The structure was positioned in a cubic box (10 Å minimum distance from box-walls) that was filled with spc216 water molecules (TIP3P). The solution was neutralized by adding 8 Cl^-^ ions and then brought to the final concentration of 150 mM NaCl. The system was then energy minimized using the steepest descent algorithm. From this structure, three independent equilibrations with position restraints on heavy atoms were launched: 500 ps in the NVT ensemble at 310 K (using the V-rescale thermostat) (32) followed by 500 ps of NPT equilibration at 350 K and 1 Bar (using the V-rescale thermostat and Parrinello-Rahman barostat) (33). Position restrains were then removed and for each equilibrated system 20 ns of molecular dynamics (in the NPT ensemble) were performed with distance restraints on the hydrogen bond lengths. The restraining potential is defined below:

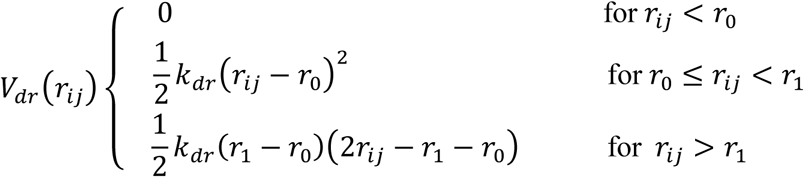

Where *r*_*ij*_ is the Euclidean distance between the H and the O atoms forming the hydrogen bond; *r*_*0*_ is the initial hydrogen bond distance (2 < *r*_*0*_ < 2.5) Å, *r*_*1*_ is an upper limit equal to 4.5 Å, and *k*_*dr*_ is set to 2·10^3^ kJ/(mol·nm^2^). The distance restraints were finally removed and three independent 100 ns trajectories were obtained in the NPT ensemble. MD simulations, together with equilibrations, were performed by employing the leap-frog integrator with time-step equal to 2 fs. Cut-off for short-range Coulomb and Van der Waals interactions was set to 12 Å and long-range electrostatics was treated using Particle Mesh Ewald. The root mean squared deviation of atomic position (RMSD) was computed on all-atoms using Gromacs 2018 while the secondary structure content was computed using VMD 1.9.2. (34). Graphs were generated with Matplotlib (35) in Python.

## ACKNOWLEDGEMENTS

The authors would like to thank the excellent technical assistance of Patricia Piñeiro. This work has received financial support from the European Union (European Regional Development Fund-ERDF); the Spanish Ministry of Industry and Competitiveness (grants BFU2017-86692-P, RTI2018-098515-B-I00, and SEV-2016-0644); the Galician Regional Government (grants ED431C 2018/04 and ED431B 2017/19); Interreg POCTEFA148/16; and Fondazione Telethon (TCP14009). E.B. is an Assistant Telethon Scientist at the Dulbecco Telethon Institute.

## AUTHOR CONTRIBUTIONS

M.M.P., V.S.P., E.B., G.S., Y.C., J.C. and J.R.R. designed the research; M.M.P. designed NMR experiments and acquired NMR spectra; M.M.P., V.S.P., Y.C. and J.R.R. interpreted NMR spectra; Y.C., P.P. and M.B. designed and performed FTIR analyses; M.D.and H.R. prepared and characterized PrP amyloid samples; S.V. prepared and characterized recPrP samples; S.V., H.E., R.L.M. and J.C. prepared infectious recPrP^Sc^ samples with methods developed by H.E. and J.C.; G.S., E.B. and J.R.R. constructed the recBVPrP^Sc^ model; G.S. performed and analyzed M.D.; R.L.M., A.I., L.F.F., J.R.R. and S.B. developed, performed and interpreted data from limited proteolysis/MS analyses of samples; M.M.P., V.S.P., Y.B. and J.R.R. wrote the paper; all authors read the final draft and made suggestions that contributed to the definitive manuscript.

## SUPPLEMENTARY MATERIAL

**Supplementary Figure 1.**
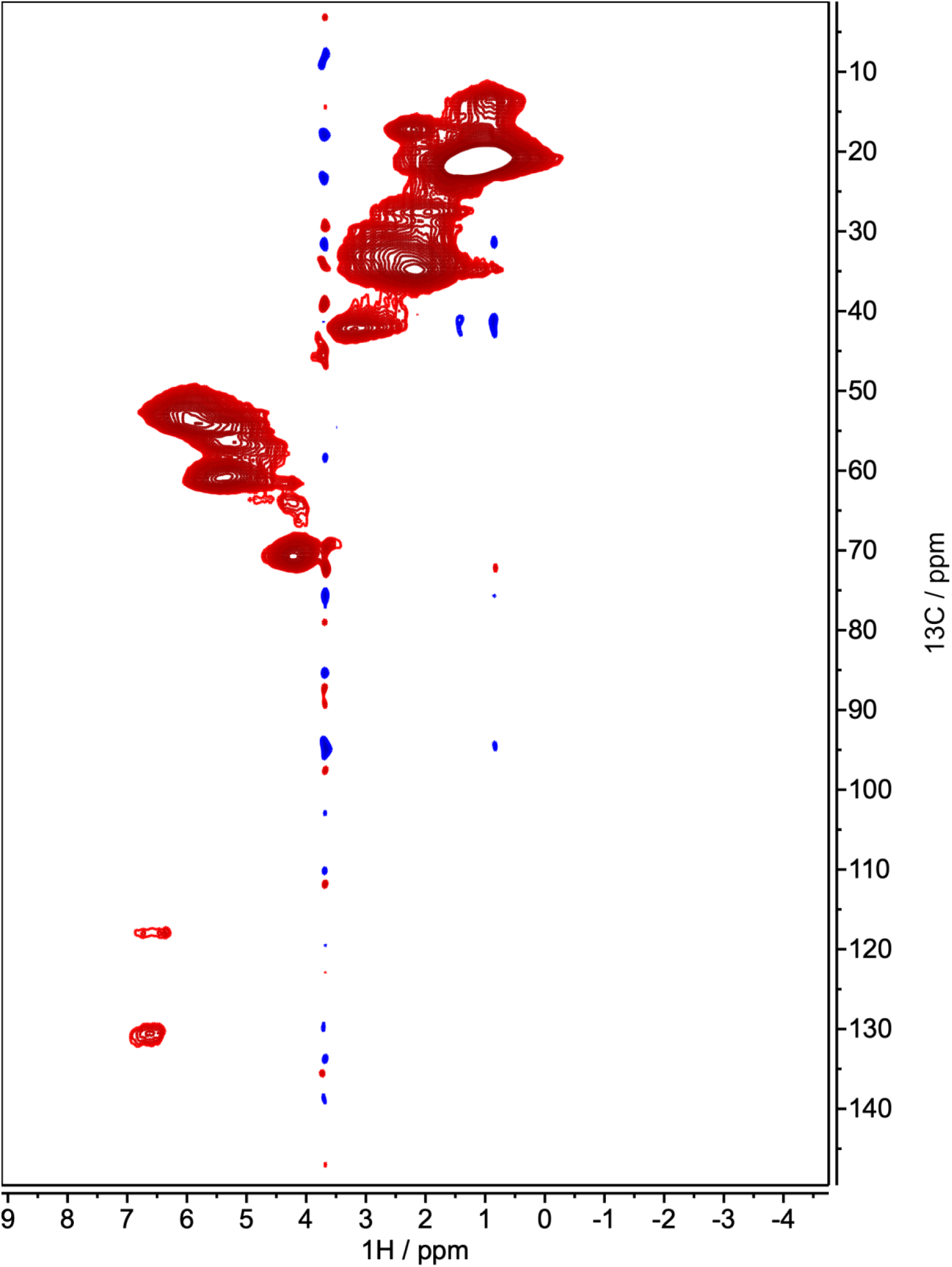
Complete 2D ^13^C-^1^H HSQC spectra of PK-treated, uniformly labelled (U-^13^C,^15^N)-BVPrP^Sc^(109I).

**Supplementary Figure 2.**
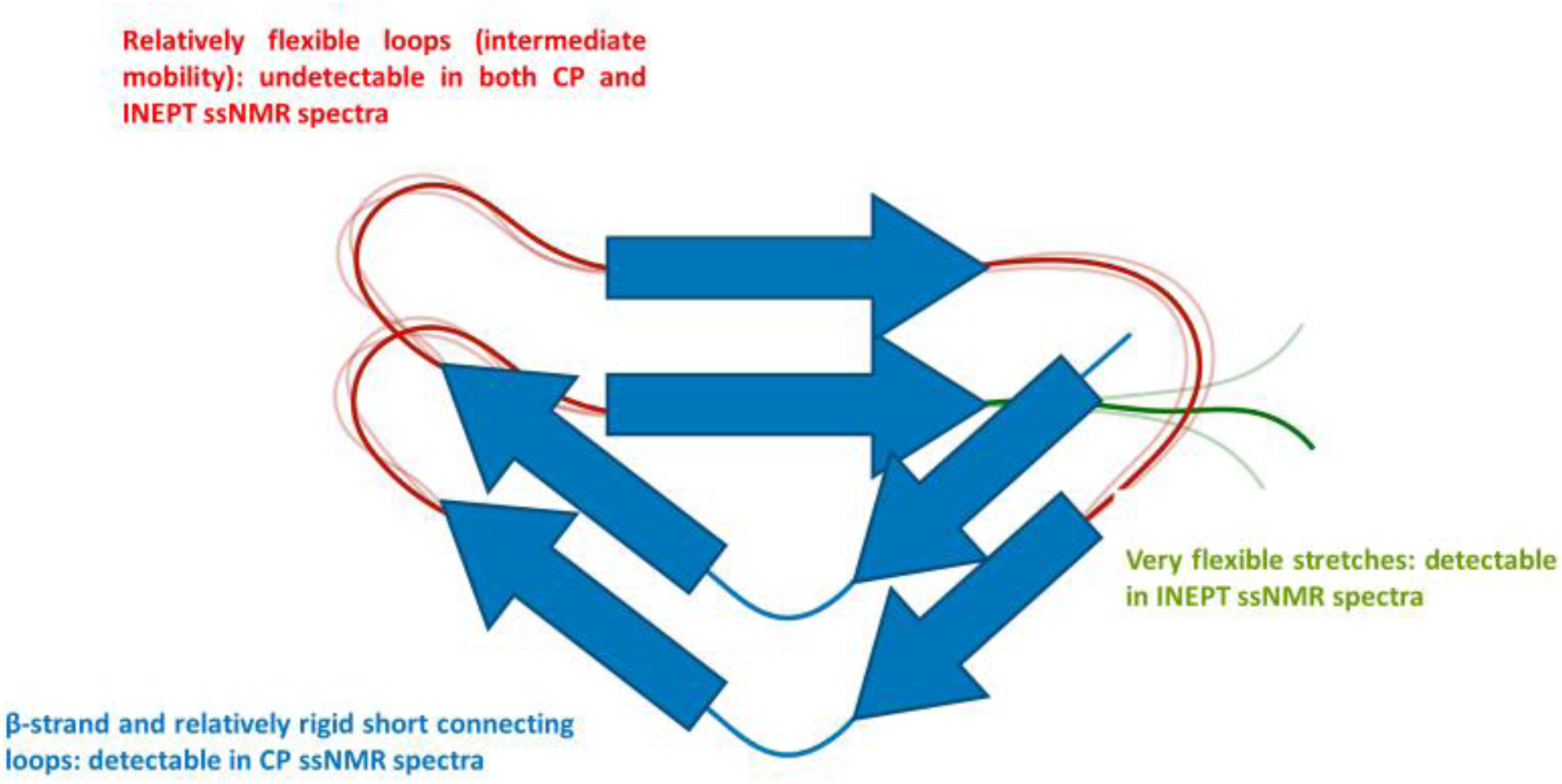
Schematic representation of the different molecular mobility regimes existing in an amyloid sample and its relationship with detectability in ssNMR spectra. Well-structured stretches have low mobility and are optimally detected in CP ssNMR spectra. Very flexible stretches (long loops or open ends) undergo fast motions even in the solid state in partially hydrated samples, which render them undetectable under standard CP ssNMR conditions; however, these residues can be detectable in INEPT-based experiments at high MAS rates. The intermediate case is that of segments with mobility at intermediate rates (i.e. with moderate flexibility); these residues may be undetectable both in CP and INEPT NMR experiments due to severe line broadening.

**Supplementary Figure 3.**
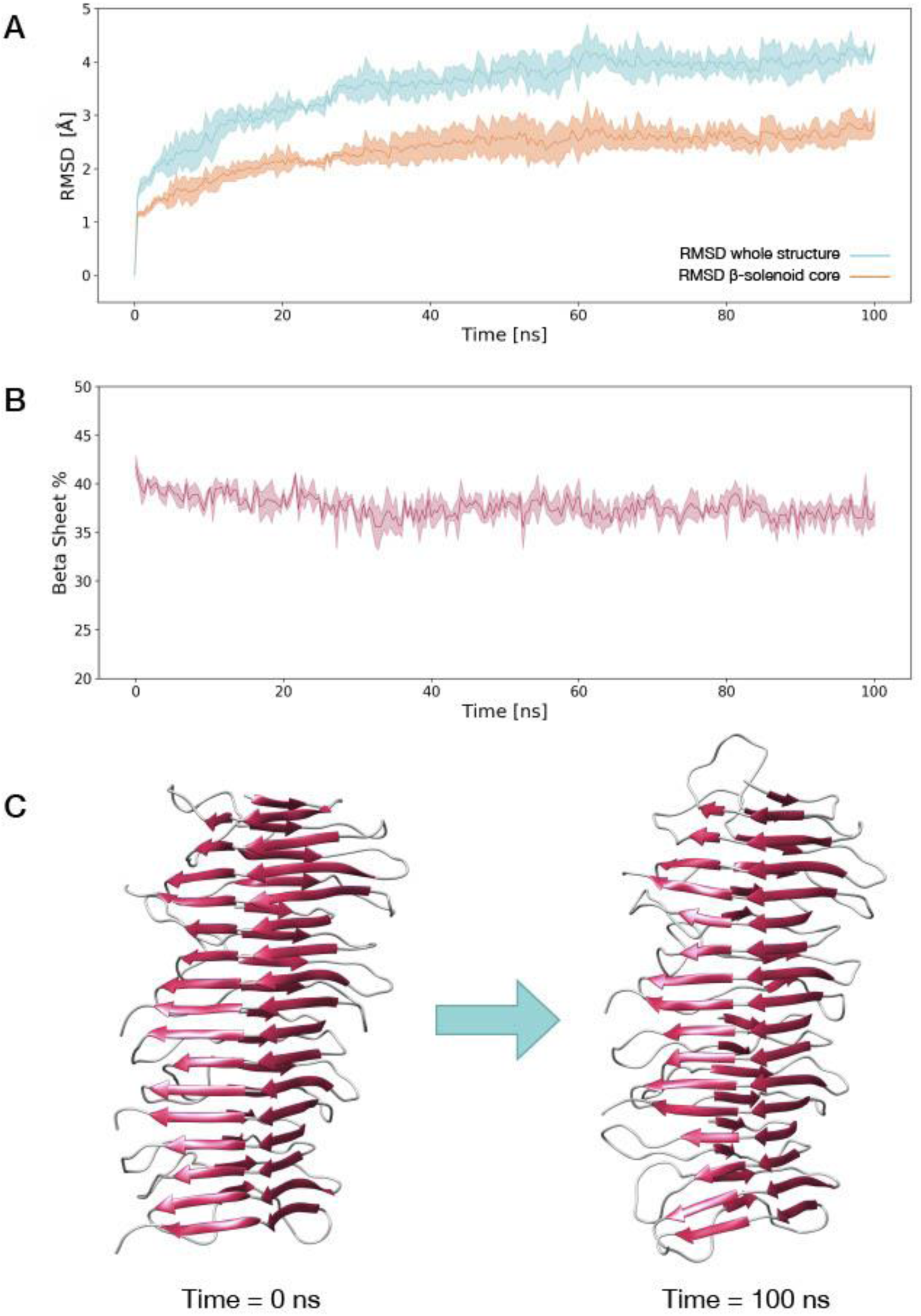
Molecular Dynamics Simulation of the BVPrP^Sc^ tetramer model. **A**) The graph shows the root mean squared deviation (RMSD) of atomic positions of the BVPrP^Sc^ tetramer model (mean of 3 simulations) accounting the entire structure (aquamarine) or the β-solenoid-core only (orange). Filled curves indicate the standard deviation. The first frame of the simulation was used as a reference. The mean RMSD in the last 10 ns corresponds to 4.0 ± 0.1 Å (mean ± standard deviation) for the entire structure and 2.7 ± 0.1 Å for the β-solenoid-core only. **B**) Percentage of β-sheet along the course of the simulation (mean of N = 3), filled curve indicates the standard deviation. The extended conformation content changes from an initial value of 39 ± 1 % (mean of the first 10 ns, ± standard deviation) to a final value of 37 ± 1% (mean of the last 10 ns, ± standard deviation). RMSD and percentage of secondary structure analysis are both consistent with the results of the simulations performed on the mouse 4RβS tetramer model reported in (4) **C**) Representative snapshots of the initial (t = 0 ns) and final (t = 100 ns) frame of the molecular dynamics simulations.

